# Plasticity of the dopaminergic phenotype and of locomotion in larval zebrafish induced by changes in brain excitability during the embryonic period

**DOI:** 10.1101/2021.07.19.452915

**Authors:** Sandrine Bataille, Hadrien Jalaber, Ingrid Colin, Damien Remy, Pierre Affaticati, Cynthia Froc, Philippe Vernier, Michaël Demarque

## Abstract

During the embryonic period, neuronal communication starts before the establishment of the synapses with alternative forms of neuronal excitability, called here Embryonic Neuronal Excitability (ENE). ENE has been shown to modulate the unfolding of development transcriptional programs but the global consequences for the developing organisms are not all understood. Here we monitored calcium transients in zebrafish embryos as a proxy for ENE to assess the efficacy of transient pharmacological treatments to either increase or decrease ENE. Increasing or decreasing ENE for 24 hours at 2 days post fertilization (dpf), at the end of the embryonic period, promoted respectively an increase or a decrease in the numbers of dopamine (DA) neurons in the telencephalon and in the olfactory bulb of zebrafish larvæ at 6 dpf. This plasticity of dopaminergic specification occurs within a stable population of vMAT2-positive cells, hence identifying an unanticipated biological marker for this reserve pool of of DA neurons that can be recruited by increasing ENE.

Modulating ENE also affected larval locomotion several days after the end of the treatments. In particular, the increase of ENE from 2 to 3 dpf promoted hyperlocomotion of larvæ at 6 dpf, reminiscent of endophenotypes reported for Attention Deficit with Hyperactivity Disorders and schizophrenia in zebrafish. These results provide a convenient framework to identify environmental factors that could disturb ENE as well as to study the molecular mechanisms linking ENE to neurotransmitter specification, with relevance to the pathogenesis of neurodevelopmental disorders.

**Significance Statement:**

- Spontaneous calcium transients, used as a proxy for Embryonic Neuronal Excitability (ENE), are detected in the forebrain of embryonic zebrafish.
- Short-term pharmacological treatments by bath application could increase or decrease ENE.
- The post-mitotic differentiation of the dopaminergic phenotype is modulated by ENE in the zebrafish forebrain.
- The plasticity of the dopaminergic specification occurs within a reserve pool of vMAT2-positive cells.
- Transient increase of ENE at the end of the embryonic period induces hyperlocomotion, a phenotype associated with ADHD and schizophrenia in this model.
- Our results open clinically relevant perspectives to study the pathogenesis of neurodevelopmental disorders in zebrafish.

## Introduction

During brain development, specific molecular components of the synaptic neuronal communication become functional before synapse formation, such as voltage sensitive ion channels and neurotransmitter receptors at the plasma membrane, and release of neurotransmitters in the extracellular space (Spitzer et al., 2002). They contribute to immature forms of cellular excitability and intercellular communications that we refer to here as Embryonic Neuronal Excitability (ENE). There is for instance a paracrine communication mediated by non-synaptic receptors activated by endogenous neurotransmitters, in the neonatal rat hippocampus, and in the mouse spinal cord (Demarque et al., 2002; Owens & Kriegstein, 2002; Scain et al., 2010). Acute changes of calcium (Ca^2+^) concentration with different spatio-temporal dynamics have also been described in differentiating neurons, either locally, in filopodia or growth cone or globally, at the level of the soma (Gomez & Spitzer, 1999; Gomez et al., 2001). The latter, we refer here as Ca^2+^transients, are sporadic, long lasting global increases of intracellular concentration of Ca^2+^ that occurs during restricted developmental windows called “critical periods”. They have been identified in the developing brain of several vertebrate species (Owens & Kriegstein, 1998; Crepel et al., 2007; Blankenship & Feller, 2010; Demarque & Spitzer, 2010; Warp et al., 2012). Changes in the incidence and frequency of Ca^2+^ transients have been shown to modulate the specification of the neurotransmitter phenotype in various populations of neurons, as is the case for dopamine (DA), in the xenopus and rat brain, with consequences on several behaviors (Dulcis & Spitzer, 2008; Dulcis et al., 2013, 2017).

DA is an evolutionary conserved monoamine involved in the neuromodulation of numerous brain functions in vertebrates, including motivational processes, executive functions and motor control (Klein et al., 2019). Accordingly, alterations of the differentiation and function of the neurons synthetizing DA contribute to the pathogenesis of several brain diseases with a neurodevelopmental origin, such as Attention Deficit Hyperactivity Disorder (ADHD) and schizophrenia (SZ) (Lange et al., 2012; Murray et al., 2017).

Further addressing the complex molecular and cellular mechanisms linking developmental excitability, dopaminergic differentiation and behavioral outputs requires an *in vivo* approach in an accessible and genetically amenable animal model with possible parallels with the human situations. The developing zebrafish *Danio rerio* fits these needs. The development of the embryo is external, which allows perturbing ENE at stages that would be *in utero* in mammalian models. The embryos are relatively small and transparent, simplifying high-resolution imaging of the brain in live or fixed preparations. Despite difference in brain organization, notably a pallium very different form the 6-layered cortex of mammals, the main neuronal systems of vertebrates are present in zebrafish and respond to psychoactive drugs (Gawel et al., 2019). In addition, the monoaminergic system has been extensively studied in zebrafish (Schweitzer & Driever, 2009; Schweitzer et al., 2012). In vertebrates, DA modulates executive functions, such as working memory and decision making, through innervation of specific regions of the telencephalon. In the zebrafish brain, most DA neurons innervating the telencephalon have their cell bodies located within the telencephalon itself (SP-DA cells) (Tay et al., 2011; Yamamoto et al., 2011). To the best of our knowledge, plasticity of the neurotransmitter phenotype in these cells has not been studied so far.

To perturb ENE in conditions close to physiological exposure and ecotoxicology, we used pharmacological treatments by bath application from 48 to 72 hours post fertilization (hpf). We then analyzed the consequences of these transient pharmacological treatments a few days later, between 6 and 7 dpf. We report quantifiable changes in the specification of the dopaminergic phenotype in SP-DA neurons. We also report induced changes in locomotion within the same time frame. These results suggest a role of ENE on the specification of the dopaminergic phenotype and motor control in the zebrafish. They also open important perspectives for this model to decipher the chain of events leading from environmental factors to the pathogenesis of brain disorders such as schizophrenia and ADHD.

## Materials and Methods

### Fish strains

All experiments were carried out in accordance with animal care guidelines provided by the French ethical committee and under the supervision of authorized investigators.

Zebrafish were raised according to standards procedures (Westerfield, 2000). Briefly, for breeding, male and female zebrafish were placed overnight, in different compartments of a tank with a grid at the bottom that allows the eggs to fall through. The next morning the separation was removed and after few minutes, the eggs were collected, rinsed and placed in a Petri dish containing embryo medium (EM). Embryos were kept at 28 C, then staged according to standard criteria. The number of animals used for each experiment is indicated in the corresponding figure legends.

Wild-type zebrafish were of AB background. The following transgenic zebrafish lines were used: Tg(hsp70l:Gal4)^kca4^; Tg(UAS:GCaMP6f;cryaa:mCherry)^icm06^; Tg(tbp:Gal4;myl7:cerulean)^f13^.

### Pharmacological treatments

Pharmacological compound, veratridine (10 μM), tetrodotoxin (TTX, 2 μM), ω conotoxin (0.08 μM), nifedipine (0.4 μM) and flunarizine (2 μM) were purchased form R&D System (UK) and prepared in water except veratridine which required dimethyl sulfoxide (DMSO) for dissolution and flunarizine that requires ethanol for dissolution. Control exposures were performed using the same concentration of DMSO in EM without the drug. Specific period and duration of applications are indicated in the corresponding figure legends. All pharmacological treatments were performed by bath application followed by three washes in EM. Embryos were randomly distributed in wells (30 embryos per well) of a 6-well plate containing 5 mL of solution (EM+DMSO or EM+drug). Embryos exposed to drugs or the control solution were observed for morphological abnormalities every day until 5 dpf. Malformations (*e*.*g*. spinal curvature, cardiac edema) were considered as experimental end-points and when detected the corresponding animals were excluded from the study.

### Calcium imaging

For Ca^2+^ imaging we measured the fluorescence of the genetically encoded Ca^2+^ sensor GCaMP6f, expressed under the control of a UAS promoter (UAS:GCaMP6f). We used two different gal4 lines to drive UAS-dependent expression of GCaMP: a (Hsp70:Gal4) line, in which the UAS transgene is expressed upon heat shock activation and a (TBP:Gal4) line, in which the TATA-box binding protein (TBP) promoter drives constitutive ubiquitous expression of the transgene.

At 24 hpf embryos of the (Hsp70:GAl4;UAS:GCaMP6f) line were exposed to a 38°C temperature for 1.5 hour. Upon heat shock activation, GCaMP6f is expressed in all the cells of the animals and remains detectable for several days.

At 2-3 dpf the embryos were paralyzed with intramuscular injections of 750μg/ml α-bungarotoxin (Life Technologies, Waltham, USA), then individually embedded in low melting agarose (Life Technologies), ventral side up, for imaging.

30 min time lapse series were acquired at 1 Hz, at a single focal plane, on an Olympus BX60 microscope (Olympus corporation, Tokyo, Japan) equipped with a 40x 0.6N water immersion objective. A non-laser spinning disk system (DSD2, ANDOR Technology, Oxford, UK) was used for illumination and image acquisition. Images were processed with Fiji. Movements of the preparation in the X/Y axis were corrected using the “Stackreg” plugin (W. Rasband, B. Dougherty). Regions of interest (ROIs) were drawn manually over individual cell bodies and the average gray level from pixels in ROIs was measured over time using the MultiMeasure plugin (Optinav, Redmond, USA). Sequential values of fluorescence were then treated in Matlab (Mathworks). Transients were defined as increase of fluorescence higher than 2.5 times the standard deviation of the baseline. The duration and amplitude of transients was calculated using the “peak” function. Incidence was scored as the number of cells generating transients divided by the estimated total number of cells in the imaged field and was expressed as a percentage. Frequency was calculated as the total number of transients in a given cell divided by the total acquisition time and was expressed as transients per hour.

### Immunohistochemistry

#### Tissue preparations

6-7 dpf zebrafish larvæ were deeply anesthetized using 0.2% Ethyl3-aminobenzoate methanesulfonate (MS222; Merck KGaA, Darmstadt, Germany) diluted in EM, then they were fixed in ice-cold 4% paraformaldehyde (PFA; Electron Microscopy Sciences, Hatfield, USA) in 1X phosphate-buffered saline (PBS; Fisher Scientific, Hillkirch, France) containing 0.1% Tween 20 % (PBST) overnight at 4 °C. Samples were dehydrated and stored in MeOH at -20°C.

#### Immunofluorescence

Immunofluorescence was performed in 2 mL microtubes. Unless specified otherwise in the protocol, incubations were performed at room temperature (RT), and thorough PBST washes were performed between each step. The samples were first incubated in 3% Hydrogen Peroxide Solution (H_2_O_2_) in Ethanol 100% (EtOH) for 30 minutes, to deactivate endogenous peroxidases. They were then successively incubated in EtOH:Xylene 1:1 without agitation for 1 hour and, at -20 °C, in EtOH:Acetone 1:2 without agitation for 20 minutes. The washes were performed between these steps were performed in EtOH. After the final wash, samples were rehydrated in PBST.

In order to unmask the antigens, samples were incubated in PBST:Tris 150 mM pH9 for 10 minutes, then in Tris 150 mM pH9 at RT for 10 minutes and at 70 °C for 30 minutes. After PBST washes, the samples were incubated in blocking buffer 1 (10% normal goat serum (NGS), 1% triton X-100, 1% tween-20, 1% DMSO, 1% Bovin Serum Albumin (BSA) in PBS 1X) for 3 hours.

Two protocols were used for primary antibody staining, adapted from published protocols (Inoue & Wittbrodt, 2011; Xavier et al., 2017; Bloch et al., 2019). For the TH antibody, the samples were incubated with the first primary antibody (mouse anti-TH, 1:250)(Yamamoto et al., 2010) in primary staining solution (1% NGS, 1% triton X-100, 1% DMSO, 1% BSA, 0,05% azide sodium in PBST) at 4°C for 7-10 days, under gentle agitation. After washing, the samples underwent a step of refixation in PFA 4% for 2 hours at RT and were washed overnight in PBST.

The samples were then incubated in blocking buffer 2 (4% NGS, 0,3% triton X-100, 0,5% DMSO in PBST) for 1 hour at RT and incubated with a first secondary antibody (anti-mouse biotinylated, 1:200)(Alunni et al., 2013) in secondary staining buffer (4%NGS, 0,1% triton X-100 in PBST) for 2,5 days at 4°C under gentle agitation.

For the revelation, we used the Vectastain ABC kit (Vector ®). Briefly, AB mix was prepared by adding 10 μl of solution A and 10 μl of solution B in 1mL PBST/1% Triton-X100. One hour after the preparation, the samples were incubated in the AB mix for 1 hour. Samples were then incubated in Tyramide-TAMRA (1:200 in PBST) for 20 minutes, then 0,012% H_2_O_2_ was added directly in the solution and the samples were incubated for an additional 50 minutes.

Before a second primary antibody incubation, the samples underwent a step of fixation in PFA 4% for 2 hours at RT and washed overnight in PBST.

For the other primary antibodies (rabbit anti-caspase3, 1:500 or chicken anti-GFP, 1:500) (Xavier et al., 2017), the samples were incubated in blocking buffer 1 for 3 hours, then were incubated with the primary antibody in primary staining solution at 4 °C for 3-4 days, under gentle agitation. After washes and refixation as above, the samples were incubated with the second secondary antibody (goat anti-chicken Alexa Fluor 488, 2 μg/ml) and DAPI 1X in PBST at 4 °C for 2,5 days under gentle agitation. Samples were then washed three times in PBST and left overnight in PBST. For observation, the brain were dissected and mounted between slides and coverslips in Vectashield solution (Vector®).

#### Image acquisition

A Leica TCS SP8 laser scanning confocal microscope with a Leica HCTL Apo × 40/1.1 w objective was used to image the specimens.

Fluorescence signal was detected through laser excitation of fluorophores at 405, 488, 552, or 638 nm and detection was performed by two internal photomultipliers. Steps in the Z-axis were fixed at 1 μm. Acquired images were adjusted for brightness and contrast using ImageJ/FIJI software.

#### Quantification of immuno-reactive cells

The R software ‘sample’ function was used to attribute a random number to each samples allowing for counting by observers blinded to treatment group. The TH- and GFP-immuno-reactive cells were counted manually from z-stacks of confocal images using the ImageJ cell counter plugin.

### Spontaneous locomotion assays

Individual larvæ were placed in a well of a 24 well plate with 2 mL EM 2 hours before recordings for habituation. The plate is then placed in a Zebrabox (Viewpoint, Lyon, France) for 10-minute recording sessions. Locomotor activity was recorded using ZebraLab software (Videotrack; ViewPoint Life Sciences, France). After a first session without threshold for 24 untreated larvæ the average speed (av_sp) of the batch was extracted from the data. For subsequent sessions, events with a speed below half the average speed (av_sp/2) were considered as inactivity. This low threshold was set in order to eliminate activity that did not contribute to the behavioural phenotype, such as small changes in orientation.

A first round of analysis was performed on the remaining events considered as swimming episodes. A second threshold was then applied, set to twice the average speed (av_sp*2). Episodes with a speed (e.s) over this high threshold were considered as fast episodes or bursts while all the episodes with a speed falling between the two thresholds were considered as normal swim episodesor “cruises”.

For each session, each animal, each category of episodes (all episodes, cruise episodes and burst episodes), 4 parameters are extracted from the recordings: the number of episodes, the total distance covered, the duration of activity, and the average speed.

## Experimental design

### Statistical analyses

Results are shown as scatterplots overlaid with box and whisker plots showing minimum (bottom whisker), maximum (top whisker), mean (cross), median (line), 1^st^ quartile (bottom of the box) and 3^rd^ quartile (top of the box), and each individual values (dots). For means comparisons, we first performed the Shapiro-Wilk test for normality. If all samples passed the normality test we then checked for equality of variance. When all samples had similar variance, we performed an ordinary ANOVA test. When differences in variances were detected, we performed Brown-Forsythe and Welch ANOVA tests (BFW ANOVA). Multiple comparisons were then performed using Dunnett’s test. If one sample or more did not pass the normality test, we used the non-parametric Kruskal-Wallis test coupled to Dunn’s test for multiple comparisons (KW). Tests and p values are in the figures legend. p < 0.05 was considered as the level for significance and are reported as follows on graphs: P < 0.05 (*), P < 0.01 (**), P < 0.001 (***).

Statistical tests were performed using either Prism (Graphpad, San Diego, USA) except for the equality of variance test, performed with the online software Brightstat (Stricker, 2008).

## Results

### Timing of experiments

To study the contribution of Embryonic Neuronal Excitability (ENE) to zebrafish larval development, we adapted the transient pharmacological treatments previously validated in xenopus (Borodinsky et al., 2004). Bath treatments were performed during the last day of embryonic development (48-72 hpf) to avoid effects on early developmental steps such as neurulation. While ENE was measured during the course of the treatments, immunohistochemical analysis and behavior tests to assess the long-term effects were measured at 6-7 dpf, several days after the end of the treatments (Figure 1A). To increase ENE we used veratridine (10 μM), which blocks the inactivation of voltage-dependent sodium channels. To decrease ENE, we used a cocktail containing TTX (2 μM), ω-Conotoxin (0.08 μM), Nifedipine (0.4 μM) and Flunarizine (2 μM) targeting respectively voltage-dependent sodium channels; N, L and T subtypes of voltage dependent Ca^2+^ channels. We refer to this cocktail as TCNF.

**Figure 1.**
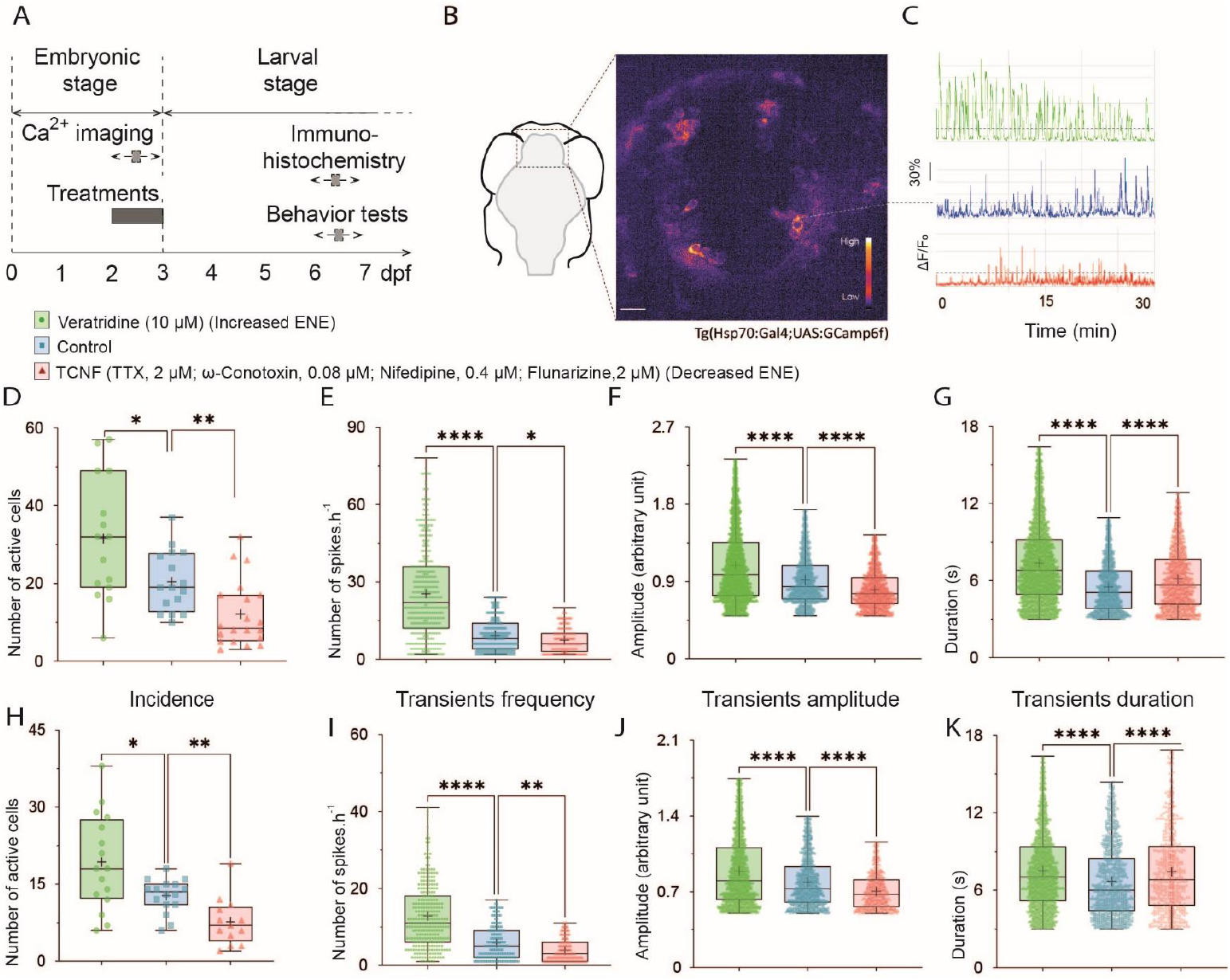
Timing of the experiments, characterization of forebrain calcium transients and their modification by bath application of pharmacological treatments. **A**. Schematic illustration of the timing of the experiments. Pharmacological treatments and calcium imaging experiments were performed from 2 to 3 dpf (embryonic period) and the immunohistochemistry and behavior experiments were performed several days later, at 6-7 dpf (larval period). **B**. Left panel, Schematic ventral view of a zebrafish brain, showing the approximate region for calcium recordings. Right panel, confocal image from a time-lapse recording of the brain of 48 hpf of Et(hsp:gal4;UAS:GCamp6f) embryos in control conditions. Fluorescence is displayed on a pseudocolor scale, the lookup table coding for the intensity scale is shown in the right corner of the image. Scale bar is 100μm. White dash circle defines an example region of interest corresponding to the cell body of a cell which change in fluorescence are displayed in C. **C**. Illustration of changes in fluorescence intensity plotted as a function of time in different experimental conditions. Representative traces from control conditions are shown, in blue, following veratridine treatment (10 μM) in green and following TCNF treatment (TTX, 2.5 μM; ω-Conotoxin, 0.1 μM; Nifedipine, 0.5 μM and Flunarizine, 2.5 μM), in red. Ca^2+^ transients were scored as changes in fluorescence more than two times higher than the SD of the baseline (dashed lines), and >3 sec in duration, calculated as the width at half-maximum. **D-K**. Boxplots showing different parameters of calcium spikes in control conditions (blue), following 10 μM veratridine treatment (green) and following TCNF treatment (red). D-G. Results obtained in the Hsp gal4 line from 5 independent experiments. H-K. Results obtained in the TBP gal4 line, from 6 independent experiments. D. Average incidence of the recorded Ca^2+^ transients, BFW ANOVA, 15<n<21 fields of view. E. Average frequency, KW test, 291<n<535 values. F. Average normalized amplitude of the recorded Ca^2+^ transients, KW test, 1258<n<4809 transients. G. Average duration of the Ca^2+^ transients, 1258<n<4809 transients. H. Average incidence of the recorded Ca^2+^ transients, BFW ANOVA, 13<n<17 fields of view. I. Average frequency, KW test, 148<n<281 values. J. Average normalized amplitude of the recorded Ca^2+^ transients, KW test, 607<n<2980 transients. K. Average duration of the Ca^2+^ transients, KW test, 607<n<2980 transients.

### Spontaneous calcium transients in the brain of zebrafish embryos

To evaluate the level of Embryonic ENE in the embryonic zebrafish brain, we used Ca^2+^ transients as a proxy. We followed the dynamics of intracellularCa^2+^ concentration using time-lapse imaging of the anterior most part of the brain of 2-3 dpf zebrafish embryos expressing the genetically encoded Ca^2+^ reporter GCaMP6f (Figure 1B). Because of the non-cell autonomous mechanism involved in the plasticity of the neurotransmitter phenotype described in xenopus (Guemez-Gamboa et al., 2014), we used ubiquitous reporters to detect changes in fluorescence in telencephalic cells of different nature including putative differentiating dopaminergic neurons, non-dopaminergic neurons and glial cells. To induce UAS-dependent GCaMP6f expression, we first used the (Hsp70:GAl4 x UAS:GCaMP6f;cry:mCherry) line, in which the transgene is expressed upon heat shock activation (see Materials and Methods and Fig 1D-G). To ensure that the observed phenotypes were not altered by the initial heat shock, we confirmed these results using a TBP:Gal4 driver, in which the TATA-box binding protein (TBP) promoter (Burket et al., 2008) drives constitutive transgene expression (Fig 1H-K).

To monitor changes in fluorescence over time, we performed time-lapse imaging of the anterior most part of the brain of 2-3 dpf zebrafish embryos for 30 minutes sessions. We analyzed the evolution of the fluorescence from selected region of interest (ROI) corresponding to individual cell bodies. We used four parameters to analyze the results, the average frequency of transients per ROI, the incidence of active ROIs in the field of view (number of ROIs displaying at least one transient during the recordings over the estimated total number of ROIs), and the duration and amplitude of individual transients (Materials and methods). In control conditions, we detected the presence of sporadic spontaneous transients in the telencephalon using both transgenic backgrounds (Figure 1D-K, blue symbols). The analyzed parameters were in the range of what has been reported in the zebrafish spinal cord, confirming that the duration of Ca^2+^ transients is shorter in zebrafish than in xenopus (Dulcis & Spitzer, 2008; Warp et al., 2012; Plazas et al., 2013).

### Global pharmacological modifications of the dynamics of calcium transients

To assess the ability of the pharmacological treatments to modify ENE, we measured their impact on Ca^2+^ signaling. We performed time-lapse recordings two to ten hours after the addition of the treatments to the embryo medium. Here again, the results obtained were similar in both genetic backgrounds. Following veratridine treatment, the four parameters analyzed were increased (Figure 1D-K, green symbols). In contrast, following TCNF treatments, three of the analyzed parameters were decreased, frequency, incidence, and amplitude while the duration was increased (Figure 1D-K, red symbols).

These results demonstrate that bath-mediated pharmacological treatments were able to change the incidence, frequency and amplitude of spontaneous Ca^2+^ transients in the embryonic zebrafish forebrain. When analyzing the late consequences of these transient treatments, we refer to the veratridine treatment as “increased ENE”, and to TCNF treatment as “decreased ENE”.

### Dopaminergic cells in the telencephalon at larval stages

Next we studied the effect of the transient perturbations on the maturation of the telencephalic dopaminergic neurons distributed as two neighboring subpopulations, in the subpallium (SP), not present in xenopus and to the best of our knowledge not studied so far and in the olfactory bulb (OB), in which a plasticity of the DA phenotype has been demonstrated in xenopus (Velázquez-Ulloa et al., 2011).

To identify dopaminergic neurons in the telencephalon of 6-7 dpf larvæ we used anatomical landmarks such as the position of brain ventricles and of large fiber bundles combined to two markers of the catecholaminergic phenotype (ie dopamine adrenaline and noradrenaline): TH, the limiting enzyme of catecholamine synthesis, and the vesicular monoamine transporter (vMAT) accumulating monoamines inside exocytotic vesicles. Indeed, in zebrafish, like in mammals, adrenaline and noradrenaline-containing neuronal soma, labelled by the same markers, are not detected anterior to the midbrain-hindbrain boundary, rather, they are restricted to neuronal subpopulations located in the locus coeruleus, the medulla oblongata and the area postrema (Ma, 1994a; b, 1997, 2003). Thus, all the catecholaminergic neurons located in front of the midbrain-hindbrain boundary are likely dopaminergic neurons.

In zebrafish, TH is encoded by two paralogous genes (TH1 and TH2). TH2, is mostly located in the caudal hypothalamus and is very rare in the nuclei studied here (Panula et al., 2010). We therefore used an anti-TH antibody that identify TH1-expressing cells (referred to as TH1^+^ cells in the remaining of the manuscript) and do not recognize TH2 (Yamamoto et al., 2011). vMAT is encoded by two different genes (vMAT1 and vMAT2) but only the latter is detected in the zebrafish brain (Puttonen et al., 2017), so we used of an anti-GFP antibody to amplify the endogenous GFP signal from cells in the Tg(Et.vMAT2:eGFP) transgenic line (referred to as vMAT2^+^ cells in the remaining of the manuscript).

In the SP, the cell bodies of vMAT2^+^ cells were distributed bilaterally along the midline, and relatively close to it (Figure 2A, cyan labelling). The major pattern of fibers distribution we could detect was that of fibers projecting first ventrally and then laterally joining the wide dopaminergic lateral longitudinal tracts.

**Figure 2.**
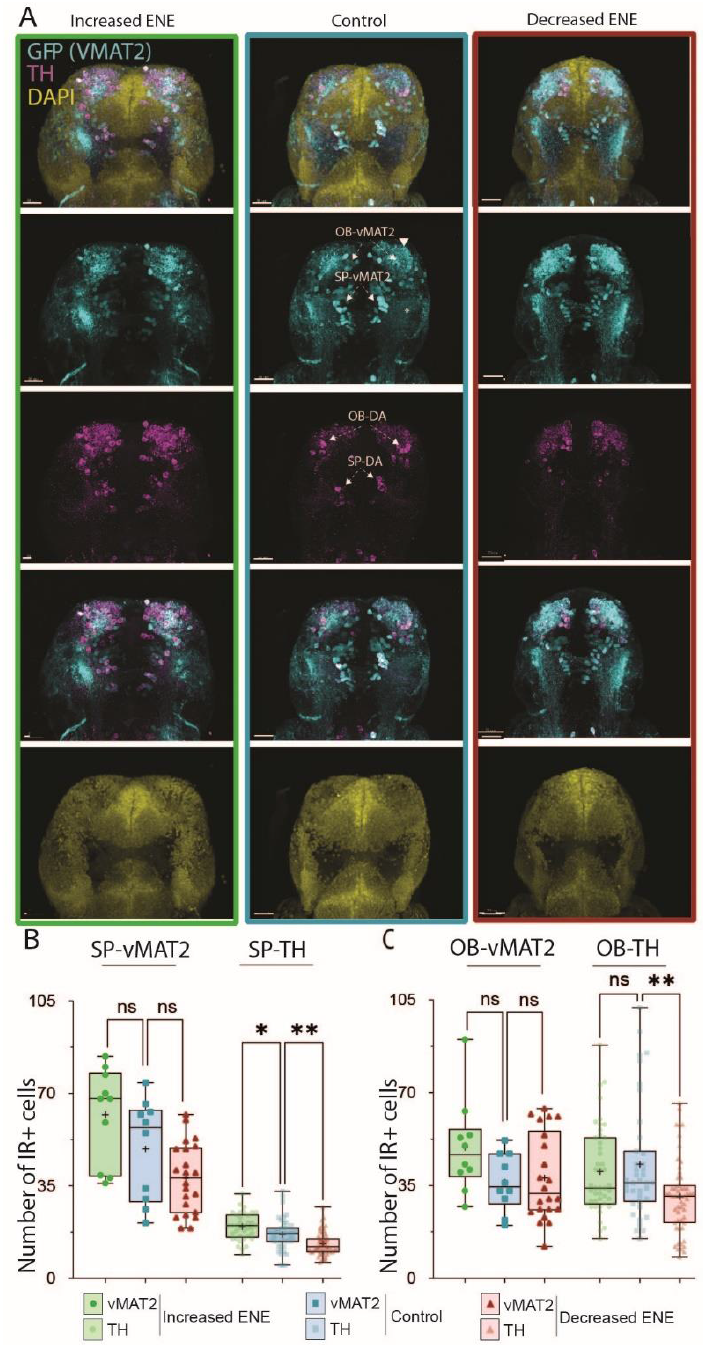
Effects of modification of ENE on the expression of dopaminergic markers in the zebrafish larval telencephalon and OB. **A**. Maximum projection of confocal z series of the brain of 6-7 dpf Et(VMAT2:eGFP) larvæ, following ENE increase (left column, boxed in green), in control conditions (middle column, boxed in blue), and following ENE decrease (right column, boxed in red). Immunostaining to TH (magenta) and GFP (cyan) and DAPI (yellow) are shown as composite image (top) as well as for each color channels. Scale bars = 50 μm. **B-C**. Boxplots showing the number of cells IR^+^ for dopaminergic markers in the SP (B) and the OB (C) from 6 independent experiments, in control conditions (blue), following ENE increase (green) and following ENE decrease (red). B. Left, number of SP-vMAT2^+^ cells, KW test, 6<n<16; right, number of SP-TH^+^ cells, KW test, 34<n<49. C. Left, number of OB-vMAT2^+^cells, KW test, 6<n<16; right, OB-TH^+^ cells, KW test, 34<n<49.

In the OB, the cell bodies of OB-vMAT2^+^ cells were distributed at the anterior end of the forebrain and they were initially projecting in an anterior direction before rapidly branching in multiple directions.

In both regions, the overall disposition of the TH1^+^ cell bodies and projections was similar as for vMAT2^+^ cells and all TH1^+^ cells were also vMAT2^+^, while some vMAT2^+^ cells were TH1^-^ (Figure 2A, magenta labelling). The population of vMAT2^+^ cells slightly exceeds that of TH1-expressing cells in the OB while the difference was more striking in the SP (Figure 2B and 2C). In the subpallium, vMAT2^+^/TH1^-^ cell bodies were mostly located at the anterior end of the cluster of vMAT2^+^/TH1^+^ cells. Such vMAT2^+^/TH^-^ cells are also detected in the subpallium of adult zebrafish (Yamamoto et al., 2011).

### Effects of modification of ENE on the expression of dopaminergic markers in the telencephalon and the OB

To assess the effects of ENE on the expression of the dopaminergic markers in the telencephalon at larval stages, we counted the number of vMAT2^+^ and TH1^+^ cells in larvæ fixed at 6-7 dpf, following the transient pharmacological treatments intended to increase or to decrease ENE.

In the SP, increasing ENE from 48 to 72hpf increased the number of TH1^+^ cells detected while decreasing ENE decreased the number of TH1^+^ cells (Figure 2A and Figure 2C). In both experimental situations, all the SP TH1^+^ cells also exhibited vMAT2 labelling. Hence, according to our identification criteria (anatomical position combined to the expression of both TH1 and vMAT2), increasing embryonic electrical activity results in more SP-DA cells in larvae, while decreasing it reduces this population. The number of vMAT2^+^ cells followed the same trend although not significantly.

In the OB, increasing ENE from 48 to 72hpf did not change the number of TH1^+^ cells detected while decreasing ENE decreased the number of TH1^+^ cells (Figure 2A and Figure 2E). As in the SP, all TH1^+^ cells exhibited vMAT2 labelling, independently of the treatments. Meanwhile, the number of vMAT2^+^ cells did not change significantly in this region (Figure 2A, Figure 2D). Overall, our results indicate that the number of dopaminergic (TH1+) neurons in the larval telencephalon is positively affected by electrical activity during the embryonic stage. Modulating ENE has a much weaker impact on the size of the monoaminergic (VMAT2+) population overall. This suggests that ENE influences the specification of monoaminergic precursors during an embryonic critical window, driving them towards the dopaminergic fate.

### Modifications of ENE has no effect on apoptosis in the forebrain

To check whether the change in the number of dopaminergic neurons following modifications of ENE are associated with programmed cell death, we counted the number of cells immunoreactive for activated caspase-3, an apoptosis marker, in the vicinity of SP and OB vMAT2 expression, at 6-7 dpf. The number of caspase-3^+^ cells was stable following the increase or the decrease of ENE (Figure 3). This result strengthens the hypothesis that the changes observed in the number of TH1^+^ cells are indeed linked to changes in neurotransmitter specification rather than changes in cell survival.

**Figure 3.**
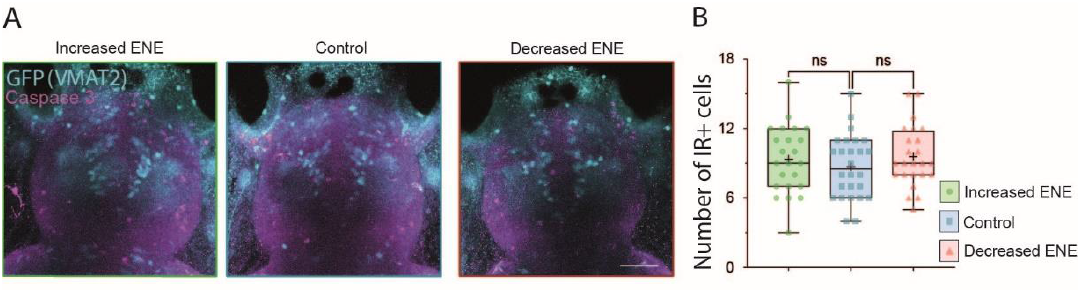
Absence of changes in programmed cell death following modifications of ENE. **A**. Maximum projection of confocal z series of the brain of 6-7 dpf Et(VMAT2:eGFP) larvæ, following ENE increase (left image, boxed in green), in control conditions (middle image, boxed in blue), and following ENE decrease (right image, boxed in red). Immunostaining to caspase-3 (magenta) and GFP (cyan) are shown as merged channels. Scale bars = 50 μm. **B**. Boxplots showing the number of the caspase-3^+^ IR cells counted in the telencephalonfollowing ENE increase (green), in control conditions (blue) and following ENE decrease (red). KW test, 23<n<26 telencephalon from 4 independent experiments.

### Effects of pharmacological treatments on the locomotion of 6-7 dpf larvæ

In order to analyze the consequence of the modifications of ENE on behaviors in zebrafish larvæ, we focused our analysis on spontaneous locomotion.

During locomotion, zebrafish larvæ display successive turn and swim bouts called episodes, interleaved with resting periods. This behavior can be analyzed by semi-automated methods, detecting 4 parameters, the number of episodes, the average duration of swimming, the distance covered and the average speed.

We compared these 4 locomotion parameters recorded at 6-7 dpf, more than 3 days after the end of ENE manipulation (Figure 1A). We recorded spontaneous locomotion over 3 consecutive 10 minutes sessions in each conditions: control, increased ENE, and decreased ENE. A representative example of traces obtained for 24 well plates in each conditions is shown in Figure 4A.

**Figure 4.**
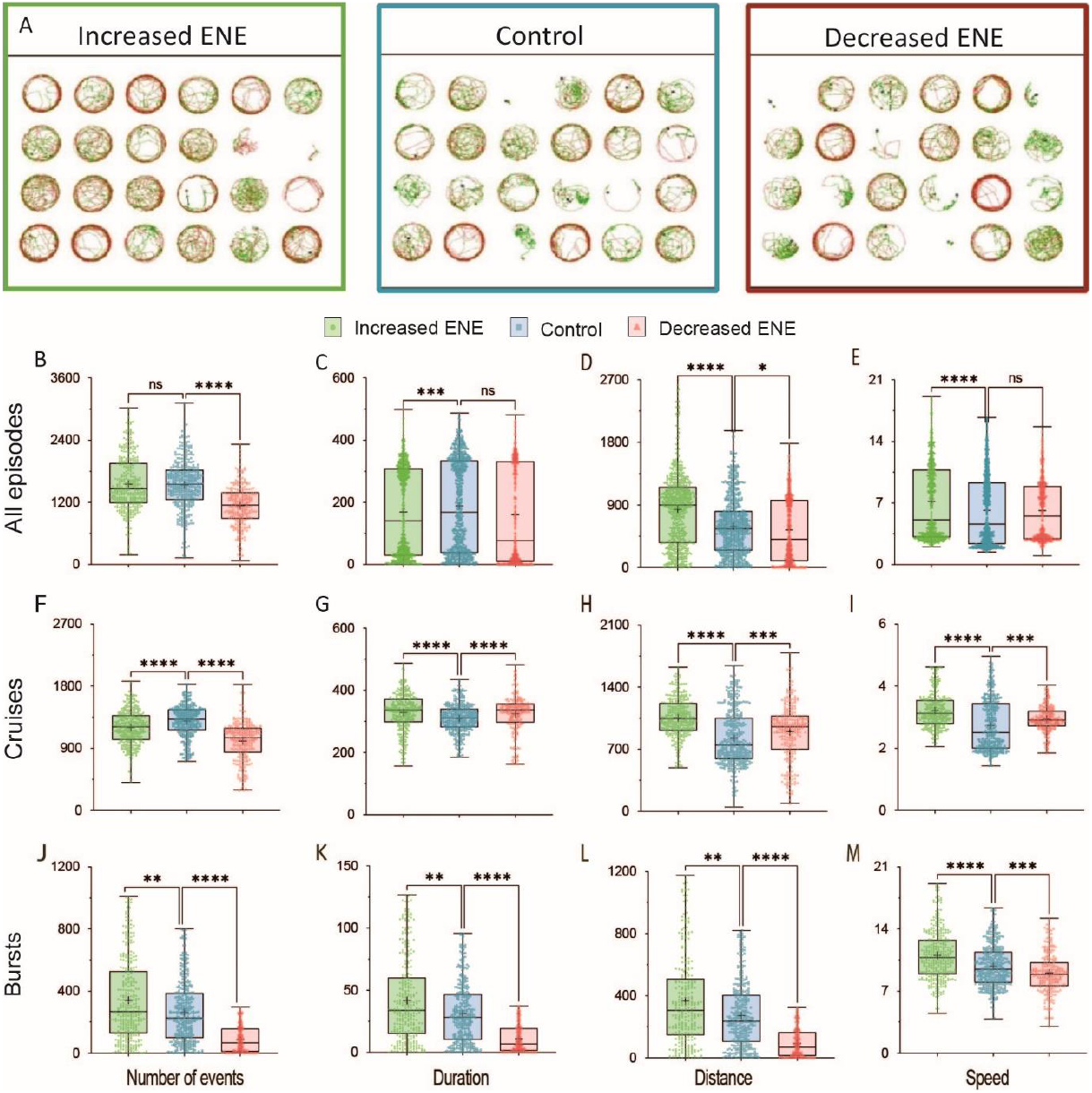
Effects of 24hrs balneation treatments during embryonic development on spontaneous swimming of zebrafish larvæ. **A**. Representative path reconstructions for 24 individual larvæ during a 10 minutes trial for three experimental conditions. We performed the experiments at 6-7 dpf when the properties of swimming episodes are relatively stable in control conditions (Extended Figure 4-1). The portion of the path corresponding to bursts episodes (shown in red) are longer following an increase of ENE, and shorter following a decrease of ENE. **B-M**. Boxplots showing the mean of different swimming parameters following ENE increase (green), in control conditions (blue), and following ENE decrease (red).. B-E Parameters when no threshold is applied for the analysis. F-I Parameters for cruises episodes. J-M. Parameters for bursts episodes. B, F, J. Number of episodes, C, G,K. Swimming duration. D, H, L. Distance covered during swimming. E, I, M. Episodes speed For all boxplots the p of normality test for at least one condition was <0.05, therefore KW test was used, 150<n<231.

Increasing ENE did not change the overall number of episodes, however it induced a decrease of the duration while the distance and speed were increased (Figure 4B-E, green symbols). Decreasing ENE decreased the number of swimming episodes and the distance, while it did not change the duration, or speed (Figure 4B-E, red symbols).

Following this observation that modifications of ENE induced changes in spontaneous larval locomotion, we analyzed further these effects by discriminating slow and fast swimming episodes, dubbed cruises and bursts, respectively (see methods and Budick & O’Malley, 2000; Kalueff et al., 2013). For cruises and bursts episodes, the same four parameters were measured for each larva tested: number of swim bouts, duration of swim, distance covered, and average speed (Figure 4 F-I and J-M respectively).

The number of cruises was reduced following either the increase or the decrease of ENE (Figure 4F). The three other parameters, duration, distance and speed, were increased in both conditions (Figure 4G-J).

By contrast, increase and decrease of ENE had opposite effects on burst episodes. Increased ENE resulted in more frequent, faster and longer bursts than in control conditions, while they were scarcer, shorter and slower after ENE decrease (Figure 4J-M).

To rule out the possibility that the differences observed following modifications of ENE could be related to a change in the speed of maturation of the locomotor network, we studied the dynamics of the 4 locomotion parameters over 4 days during the larval development in control conditions (Supplemental Figure 1). We tracked the position of individual larvæ during 3 rounds of 10-minute recordings at 4, 5, 6 and 7 dpf. The recorded parameters reflected a gradual increase of spontaneous locomotion from 4 to 6 dpf and a relative stability between 6 and 7dpf. From 4 to 6 dpf, the number of episodes, the duration of swimming and distance covered increases, both for cruises and bursts episodes. In contrast, between 6 and 7 dpf, the recorded parameters were stable both for cruises and bursts episodes. Based on these results, we concluded that differences observed at 6-7 dpf, would be related neither to a delay nor to an acceleration of the maturation of the locomotor network since the timing of recording correspond to a relative stable period for swimming parameters.

These results showed that the consequences of perturbing the embryonic electrical excitability are different on the cruise and on burst episodes, suggesting that the underlying neural networks of these swimming modes are not the same and/or that they are not regulated in the same way following changes in ENE.

### Washout kinetics of pharmacological treatments assessed using locomotion parameters at 6-7 dpf and calcium signaling at 2-3 dpf embryos

To exclude a direct contribution of the treatments applied in the embryonic period on the parameters measured in the larval period, we assessed the reversibility of the treatments by following the time course of the effects of acute pharmacological treatments in two sets of experiments.

First, on 5dpf larvae, we performed 10-minute recording sessions before application of the treatments, then +2 hours after the application of the treatments, and lastly 24 hours after washing out the drugs. We analyzed burst episodes. Acute treatments had overall similar effects as the one recorded after treatments performed in 2-3 dpf embryos. Veratridine application induced an increase of number, duration, distance, and speed of bursts, while T.C.N.F application led to a decrease of number, duration and distance of bursts, only the average speed remaining unchanged (Figure 5A-D, left boxplots). By the next day, all the effects of the treatments on bursts episodes had faded away demonstrating the washout properties of the pharmacological agents used (Figure 5A-D, right boxplots).

**Figure 5.**
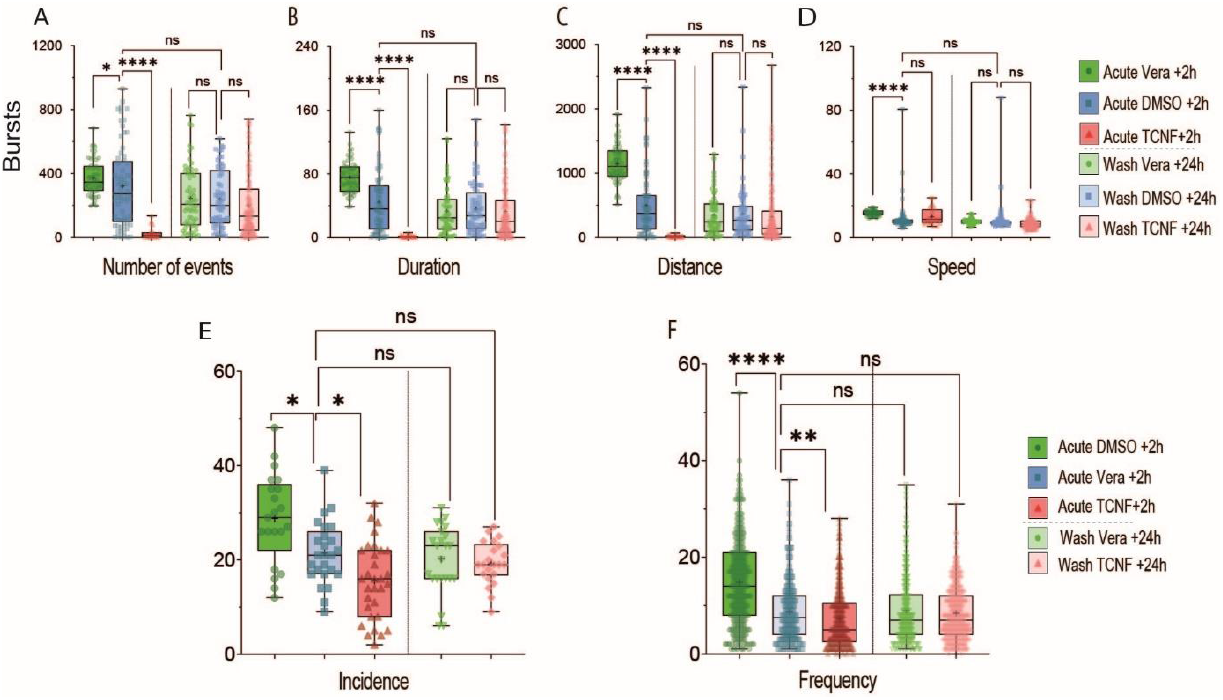
Washout kinetics of the pharmacological treatments in the larval and embryonic zebrafish brain. **A-D**. Boxplots showing the quantification of different swimming parameters for bursts episodes from 5 dpf larvae, following acute application of veratridine (green), in control conditions (blue), following acute application of TCNF (red) and after 20 to 24 hours of wash for each treatment (light boxplots). For all boxplots the p of normality test for at least one condition was >0.05, therefore KW test was used, 28<n<133 fish from 3 independent experiments. **E-F**. Boxplots showing parameters of Ca^2+^ transients in the telencephalon of 2 dpf embryos, following acute veratridine application (green), in control conditions (blue), following acute TCNF application (red) and following 6 to 10 hours of wash for each treatment (light green and light red boxplots). E Incidence of Ca^2+^ transients. BFW ANOVA test. F Frequency of Ca^2+^ transients, KW test. 258<n<318 values from 3 independent experiments.

Second, we recorded the washout kinetics of pharmacological treatments on Ca^2+^ signals recorded in 2-3 dpf embryos. We applied the drugs at 48 hpf, as before. We recorded 20-minute sessions between 2 and 4 h after the beginning of the applications, then washed the treatments, waited for 6 to 10hrs and perform new 20 minutes sessions (Figure 5 E and F). As described in Figure 1, after 2 hours of treatment, incidence and frequency of Ca^2+^ transients was increased by veratridine and reduced by TCNF (Figure 5 E and F, left boxplots). For both treatments, the values measured after the washing time (or washout) were back to control values (Figure 5 E and F, right boxplots).

These results indicate that washout of veratridine or TCNF results in rapid loss of direct effect of the drugs, in a few hours at most. Thus the timing of our analysis of larval locomotion, 3 days after ENE manipulation, is far beyond the time required for the disappearance of the acute effect of the drugs. Hence, our results support the existence of a long-term effect related to changes in excitability during the embryonic period in the zebrafish.

## Discussion

### Summary

Immature forms of excitability, here referred to as Embryonic Neuronal Excitability (ENE), include *Ca*^*2+*^ transients and activation of non-synaptic receptors to neurotransmitters and contribute to modulate neuronal differentiation. Our results show that the developing zebrafish is a suitable model to manipulate ENE using pharmacological treatments and to study the consequences of these manipulations on the plasticity of neuromodulator systems and on the behavior. Indeed, we report that ENE exerts a positive regulatory effect on the specification of the dopaminergic phenotype by increasing the number of dopaminergic neurons in the subpallium and on the properties of episodes of high speed swimming.

### Bath application of pharmacological treatments and ENE

For the present study, zebrafish embryos were exposed to pharmacological compounds by bath application. This methodological approach allows performing global and transient drugs applications, relevant to natural situations where embryos can be exposed to various biologically active or pollutants. Zebrafish is highly suitable for such questions because of its external development, in an egg. In addition, our washout experiments fit well with data in the literature indicating that the blood brain barrier starts to form around 72hpf but is not fully mature until 10 dpf (Fleming et al., 2013), allowing the diffusion of drugs to neuronal tissues during treatments and their subsequent removal upon washing at embryonic stages.

We focused on Ca^2+^ transients, one of the main forms of immature excitability, to assess the effectiveness of the treatments, but other intercellular communication processes such as the paracrine activation of GABA and glutamate receptors by endogenous transmitters are also likely perturbed by the treatments. Further studies are required to decipher the specific contribution of each of these mechanisms to the cellular and behavioral effects we reported here.

### Calcium transients in the zebrafish subpallium

One reason to focus on Ca^2+^ transients is because they are conveniently measured by Ca^2+^ imaging. In addition, they have been previously involved in neurotransmitter phenotype specification in other models of neuronal plasticity. For example, in the developing xenopus spinal cord, these Ca^2+^ transients induce release of brain-derived neuroptrophic factor (BDNF) which result in non cell-autonomous phenotype switching (Guemez-Gamboa et al., 2014). We chose to drive the expression of GCaMP ubiquitously, this ensured we would not miss relevant cell population at the selected period of recordings. The next step would thus be to use promoters of different cellular subpopulations, in particular differentiation markers of the DAergic lineage to identify the differentiation status of the cells where the Ca^2+^ transients are detected. It would also allow to see whether there are different dynamics of Ca^2+^ transients in different subpopulations of DA neuron precursors.

Ca^2+^ transients were already reported in the spinal cord of zebrafish embryos around 24 hpf (Warp et al., 2012; Plazas et al., 2013). Here, recordings were performed in the subpallium and olfactory bulb of embryonic zebrafish around 48 hpf, a time at which the transition toward synaptic network activity has already occurred in the spinal cord (Warp et al., 2012). These observations are in accordance with the existence of an postero-anterior gradient of neuronal maturation, similar to what was described in xenopus (Papalopulu & Kintner, 1996).

The frequency and duration of Ca^2+^ transients we measured in the subpallium were similar to what has been described in the zebrafish spinal cord (Plazas et al., 2013). This duration (3.3+/-1.6 s) is overall shorter than what has been reported in Xenopus (7.5+-1.0 s)(Gu et al., 1994). The basis of these interspecies differences are not known. One hypothesis is an influence of temperature since the two species do not develop at the same temperature and temperature has been shown to influence the development in xenopus (Spencer et al., 2019) but also more generally during evolution (Jaksic et al., 2020). It suggests that the duration of the transients is not necessarily the pertinent signal to trigger the effect on neuronal differentiation, a proposition further supported by the results of the veratridine and TCNF treatments reported here, which led to diametrically opposite effects on DA specification and on high speed locomotion whereas the duration of individual transients was increased in both cases. Frequency and possibly amplitude of transients appear to be more relevant candidate specification triggers.

### Potential mechanisms linking changes in ENE and plasticity of the neurotransmitter phenotype

Several mechanisms underlying the contribution of activity-dependent processes in the choice of neurotransmitter phenotype have been proposed. For neurons in the dorsal embryonic spinal cord of Xenopus tropicalis, a direct link between endogenous Ca^2+^ transients and an intrinsic genetic pathway has been shown to influence the neurotransmitter choice. Ca^2+^ signals increase the phosphorylation of the transcription factor c-Jun, in turn phosphorylated c-Jun drives the expression of tlx3, another transcription factor, which favors the GABAergic fate over the glutamatergic fate contributing to an homeostatic plasticity of the neurotransmitter phenotype specification (Marek et al., 2010).

A list of transcription factors involved in the specification of the dopaminergic phenotype expressed in dopaminergic neurons or in their vicinity in the zebrafish brain at 96 hpf is available (Filippi et al., 2012). Some may have an expression sensitive to the level of activity providing critical decision points in genetic networks for neurotransmitter specification. To test this hypothesis, the sensitivity to activity of the expression of several of these candidates will be tested in the future.

Indirect mechanisms may also be involved. Ca^2+^ transients have been shown to induce the release of BDNF which in turn triggers the activation of the TrkB/MAPK signaling cascade modulating the expression of transcription factors involved in neurotransmitter fate selection (Guemez-Gamboa et al., 2014).

In ecotoxicology exposure to modulators of activity both mechanisms are likely activated and further analysis are required to disentangle their respective contribution to the phenotypes reported here.

### Critical period for the effect of ENE perturbations

We observed an effect of the pharmacological treatments performed during the embryonic period on spontaneous locomotion several days after the end of the treatments. In contrast, no effects were observed 24 hours after treatments performed at 5 dpf in the zebrafish larvæ. These long-lasting effects of treatments specifically during the embryonic period are in line with the existence of a “critical period” during development, more sensitive to homeostatic perturbations, and promoting a significant phenotypic plasticity in immature neurons. Interestingly, dopaminergic systems have multiple functional roles and are particularly prone to plasticity events (Collo et al., 2014; Macedo-Lima & Remage-Healey, 2021), suggesting that these systems might be a key factor for adaptability of the animals to environmental changes. Whether it is a cause, a consequence or a simple correlation of their conservation throughout animal evolution is still an open question.

### Identification of a biomarker for DA reserve pool neurons

In the SP, the effect of ENE perturbations on the DA phenotype analyzed by the number of TH1^+^ cells was in agreement with the homeostatic rule described in xenopus (Spitzer, 2012). Indeed, DA phenotype was enhanced by increased excitability, and decreased by decreased excitability, as expected from an overall inhibitory neurotransmitter. In the OB, we observed a similar decrease of the DA phenotype following decreased excitability but no increase following increased activity. This unexpected result could be due to a number of TH1^+^ cells already set at a maximum in basal conditions in the OB as suggested by the high ratio of vMAT2^+^/TH^+^ observed in this region compare to SP in control conditions.

The relative stability of the number of vMAT2^+^ cells, together with no change in caspase 3-labelled cells upon treatments, suggest that modulation of excitability did not affect cell death or proliferation, but rather influenced the commitment of precursors of dopaminergic neurons. This plasticity of the DA phenotype is likely to occur within the pool of vMAT2^+^ cells, since the number of vMAT2^+^/TH^+^ was increased with increased excitability. Therefore, the vMAT2^+^/TH^-^ cells could be a reserve pool of cells primed to become dopamine neurons when plasticity-triggering events occur. The expression of the vesicular transporter in a reserve pool of cells, might have a functional advantage in term of response to plasticity-triggering events, limiting the number of factors to be changed for reaching a fully functional dopamine phenotype.

### Behavioral consequences of ENE

Perturbations of ENE had also behavioral consequences in zebrafish larvæ, modifying spontaneous locomotion. For burst episodes, the initiation of movement is likely to be the prime parameter modified following ENE perturbations, fitting with a contribution of dopamine. Determining whether a direct activation of dopaminergic neurons in the subpallium could stimulate bursts using optogenetic tools is an open perspective for the future. Since hyperlocomotion is an endophenotype related to ADHD and schizophrenia in zebrafish (Blin et al., 2008; Lange et al., 2018), it would be interesting to test whether other behaviors related to ADHD and schizophrenia such as prepulse inhibition are also affected following alterations of ENE.

### Potential mechanisms linking changes in ENE and changes in locomotion

Links between movement control and the neuromodulator effect of dopamine are widely described. In zebrafish, it has been involved in the initiation of movement (Thirumalai & Cline, 2008), via direct projection from dopaminergic cells in the diencephalon (Lambert et al., 2012). Dopaminergic cells in the hypothalamus also modulate locomotion (McPherson et al., 2016). The global perturbations we used likely induced plasticity mechanisms beyond the dopaminergic cells we analyzed in this study. Changes within the spinal networks itself, and within non-dopaminergic modulatory networks probably occurred as well, preventing us to conclude about the ties that may link the number of TelDA neurons and the level of spontaneous locomotion, the two phenotypes we report here.

### Potential involvement of changes of ENE in the pathogenesis of brain disease

Results from epidemiological studies and experiments in rodents further suggest that a range of functional and behavioral abnormalities observed in the mature system result from early alterations of monoaminergic neurons differentiation (Money & Stanwood, 2013; Suri et al., 2014). Besides genetic factors, environmental factors such as malnutrition, stress, and drugs exposure during the embryonic period could also contribute to the appearance of the pathologies. The data presented here point to an implication of ENE in the regulation of dopaminergic differentiation in the developing brain, suggesting that ENE could act as an intermediate between environmental factors and the molecular changes leading to the alteration of dopamine-related behaviors.

According to the ‘Developmental Origins of Health and Disease (DOHaD) hypothesis, transient exposure to perturbations during the development could lead to emergence of disease in young or adult individuals (Mandy & Nyirenda, 2018). The developing zebrafish provide an interesting model in which to test this hypothesis, using for instance transcriptomic analysis to identify genes differentially expressed in vMAT2 cells following modifications of environmental factors and of ENE for instance. The results would help decipher the cellular and molecular mechanisms linking exposure to external challenges with the subsequent changes in ENE, maturation of the dopaminergic systems and their behavioral outputs, with potential relevance to phenotypes related to developmental brain disorders such as ADHD and schizophrenia.

## Acknowledgments

The authors thank Kei Yamamoto for scientific inputs, TEFOR Paris Saclay (TPS) for technical support.

## Figure legends

**Extended Figure 4-1, related to Figure 4.**
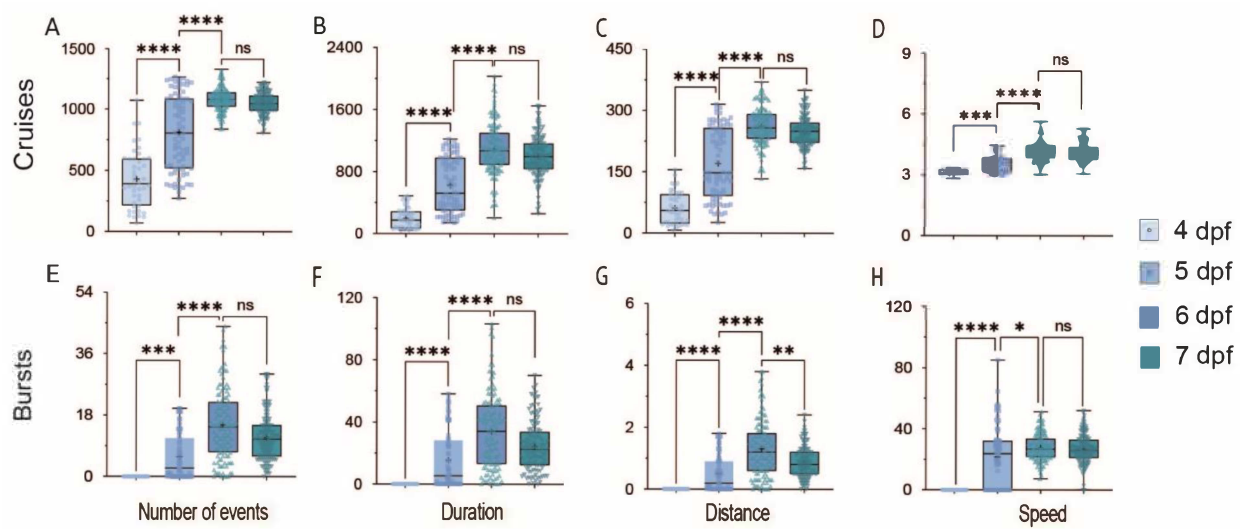
Stability of the locomotion parameters between 6 and 7 dpf. **A-H**. Boxplots showing the results for the 4 swimming parameters measured in control conditions at 4 developmental stage 4, 5, 6 and 7 dpf (coded in progressively darker blue). A,D. Parameters for cruises episodes. E-H. Parameters for bursts episodes. A,E. Number of episodes, B,F. Swimming duration. C,G. Distance covered during swimming. D,H. Episodes speed For all boxplots the p of normality test for at least one condition was >0.05, therefore KW test was used. At 4dpf, n=44, at 5dpf, n=76, at 6dpf, n= 135 and at 7 dpf, n=138, from 2 independent experiments.

